# Pathway-based clustering identifies two subtypes of cancer-associated fibroblasts associated with distinct molecular and clinical features in pancreatic ductal carcinoma

**DOI:** 10.1101/2024.12.17.628647

**Authors:** Hongjing Ai, Rongfang Nie, Xiaosheng Wang

## Abstract

Existing single-cell clustering methods are based on gene expressions that are susceptible to dropout events in single-cell RNA sequencing (scRNA-seq) data. To overcome this limitation, we proposed a pathway-based clustering method for single cells (scPathClus). scPathClus first transforms single-cell gene expression matrix into pathway enrichment matrix and generates its latent feature matrix. Based on the latent feature matrix, scPathClus clusters single cells using the method of community detection. Applying scPathClus to PDAC scRNA-seq datasets, we identified two types of cancer-associated fibroblasts (CAFs), termed csCAFs and gapCAFs, which highly expressed complement system and gap junction-related pathways, respectively. Spatial transcriptome analysis revealed that gapCAFs and csCAFs are located at cancer and non-cancer regions, respectively. Pseudotime analysis suggest a potential differentiation trajectory from csCAFs to gapCAFs. Bulk transcriptome analysis showed that gapCAFs-enriched tumors are more endowed with tumor-promoting characteristics and worse clinical outcomes, while csCAFs-enriched tumors confront stronger antitumor immune responses. Compared to established CAF subtyping methods, this method displays better prognostic relevance.

## Introduction

Pancreatic ductal adenocarcinoma (PDAC) is a highly malignant disease with an average 5-year survival rate less than 10% [1]. Its poor prognosis is attributed to late detection and early metastases that make most PDAC patients unsuitable for surgery at diagnosis [2]. A hallmark of PDAC is desmoplasia, in which fibrotic stroma constitutes a major fraction of the tumor microenvironment (TME) [3]. Fibrotic stroma may impede drug delivery, hamper immune cell access, and promote resistance to cytotoxic therapies [4]. As an abundant cell type in the TME, cancer-associated fibroblasts (CAFs) are highly versatile, plastic, and resilient [5]. Though CAFs are a promising target for anti-tumor interventions [6], CAFs-related clinical trials are hampered by the lack of specific markers as well as the high heterogeneity among CAF subtypes [5]. In PDAC, the functions of CAFs are more complex than a homogeneous tumor-promoting phenotype [7, 8]. Thus, a better understanding of the diversity of CAFs may improve personalized prognosis and treatment for PDAC patients.

Recently, the advancement of single-cell RNA sequencing (scRNA-seq) technique has enabled the identification of CAF subpopulations based on the expression of related marker genes, leading to a deep understanding of the CAF diversity in PDAC and other caner types. For example, myofibroblasts (myCAFs) express the marker genes *ACTA2* and *TAGLN* and play a tumor-suppressive role [9–11]; inflammatory fibroblasts (iCAFs) express the marker genes *IL6* and *CXCL12*, which have a tumor-promoting function through the production of inflammatory cytokines to attract immunosuppressive cells [9–11]; and antigen-presenting fibroblasts (apCAFs) play a role of immunomodulation in antigen presentation [9]. However, these definitions of CAF subpopulations based on the expression of marker genes are ambiguous. For example, the actual roles of apCAFs could not be determined as the markers of apCAFs were also overrepresented in other cell types, such as macrophages and B cells [12, 13]. It suggests that conventional single-cell clustering methods based on gene expressions have certain limitations in identifying cell subpopulations with different functions. The innate deficiencies of scRNA-seq data, such as noisy and sparse gene expressions, could be responsible for the limitation of the methods identifying cell subpopulations based on single-cell gene expressions [14]. To overcome the limitation, using pathway information in clustering analysis may improve the stability and robustness of results as the pathway expression-based clustering may remedy the instability caused by abnormality or outliers of single genes’ expression values [15–17].

In this study, we designed a pathway-based clustering method for single cells (scPathClus). We applied scPathClus to identify CAF subpopulations with different pathway activity and biological functions in PDAC. This method identified two CAF subpopulations showing different pathway activity. Furthermore, we characterized both CAF subpopulations by analyzing cell-cell communications, spatial transcriptomic data and bulk transcriptomic data, and explored their potential in precise classification of PDAC. Our data would provide novel insights into the biology of CAFs in PDAC as well as potential clinical implications for PDAC diagnosis, prognosis, and treatment.

## Methods

### Materials

From public databases, we downloaded two PDAC scRNA-seq datasets to perform pathway-based single-cell clustering, including an annotated data (PRJCA001063 [18]) from zenodo (https://zenodo.org/record/3969339) and another dataset (GSE155698 [19]) from the NCBI Gene Expression Omnibus (GEO) (https://www.ncbi.nlm.nih.gov/geo/query/acc.cgi?acc=GSE155698). Besides, we downloaded a microarray-based spatial transcriptome dataset of PDAC with matched scRNA-seq data [20] from GEO (https://www.ncbi.nlm.nih.gov/geo/query/acc.cgi?acc=GSE111672). The RNA-seq gene expression (count level) and clinical data for the TCGA pancreatic adenocarcinoma (PAAD) cohort were downloaded from the University of California Santa Cruz (UCSC) Xena platform (https://xenabrowser.net/datapages/), from which the data of 146 PDAC patients were extracted based on histological diagnosis information. In addition, we downloaded two bulk RNA-seq datasets: the PACA-AU cohort from the International Cancer Genome Consortium (ICGC) database (https://dcc.icgc.org/releases/current/Projects), and the CPTAC-PDAC cohort from the LinkedOmics portal (http://linkedomics.org/login.php).

In total, we analyzed 13,259 pathways or gene sets. Among them, 340 human KEGG pathways [21] were downloaded using the R package “KEGGREST.” The other pathways were downloaded from the Molecular Signatures Database (MSigDB) (https://www.gsea-msigdb.org/gsea/msigdb/human/collections.jsp#H), which includes 10,532 gene ontology (GO) [22] gene sets, 733 pathway gene sets from the WikiPathways, and 1,654 pathway gene sets from the Reactome database [23].

### Overview of the scPathClus algorithm

The scPathClus algorithm first uses UCell [24], a robust single-cell gene set scoring algorithm, to calculate the gene set enrichment scores for single cells and obtained the matrix 𝑍𝑍 (𝑘𝑘 × 𝑛𝑛), with the input of the single-cell gene expression matrix 𝑋𝑋 (𝑚𝑚 × 𝑛𝑛). Here *k*, *n* and *m* denote the number of gene sets, single cells and genes, respectively. Next, based on *Z*, scPathClus constructs an autoencoder (AE) to generate the latent feature matrix 𝑍𝑍^′^ (64 × 𝑛𝑛). The AE is composed of five layers: input layer, three hidden layers, and output layer. The number of nodes at each layer are *k*, 1,024, 64, 1,024, and *k*, respectively. The ReLU function and the sigmoid function are the activation function in the hidden layers and the output layer, respectively. scPathClus uses the Adam optimization algorithm with a learning rate of 0.00001 to update network weights iterative in the AE. The mean squared error (MSE) is utilized to evaluate the AE’s performance. Besides, the batch size and the epoch number are set as 64 and 50, respectively. scPathClus builts the AE with keras, a deep learning application programming interface (API) that runs on the python machine learning platform TensorFlow. After obtaining the latent feature matrix 𝑍𝑍^′^ by the AE, scPathClus utilizes the scalable toolkit scanpy [25] to analyze 𝑍𝑍^′^. This procedure produces the neighborhood graph based on 𝑍𝑍^′^and clusters single cells using the Leiden method of community detection. An illustration of the scPathClus algorithm is shown in Figure 1.

**Figure 1.**
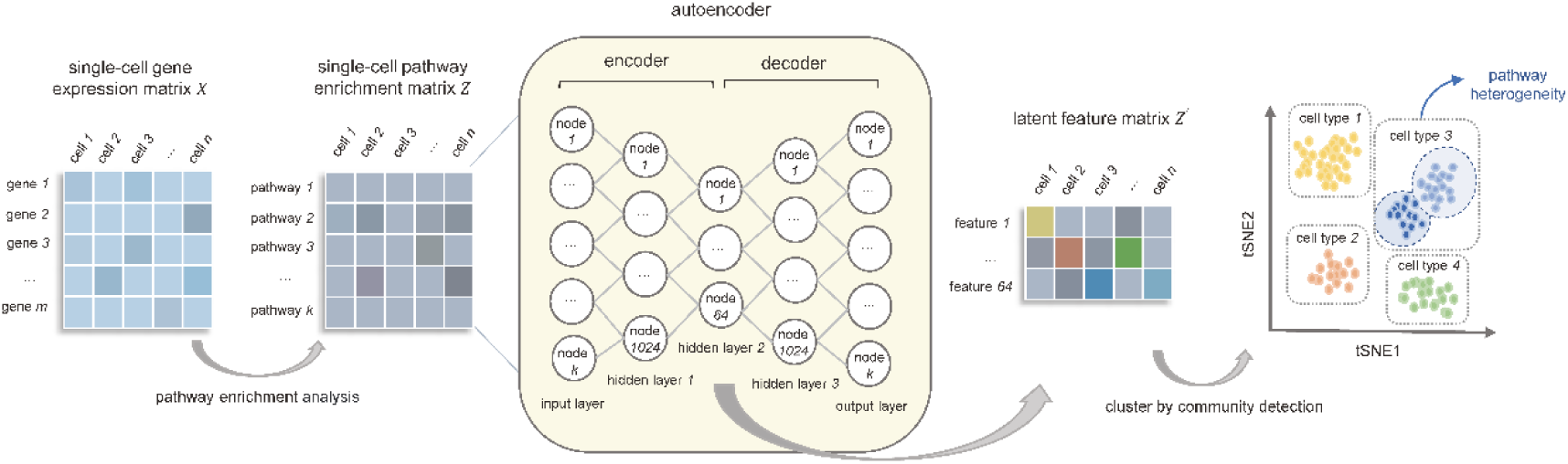
Illustration of the scPathClus algorithm.

The codes for the scPathClus algorithm are available at the website: https://github.com/WangX-Lab/scPathClus, under a GNU GPL open-source license.

### Cell-cell communication

To identify cross-talks between CAFs and tumor cells, we used the R package “CellChat” [26] to explore the secreted signaling in the CellChatDB database.

### Trajectory inference of single cells

We utilized the R package “Monocle2” [27] to build single-cell trajectories and performed the pseudotime analysis of the single cells in the CAF subtypes.

### Spatial transcriptome analysis

For each tumor spot, we used the UCell algorithm [24] to calculate gene set enrichment scores and utilized the CARD algorithm [28] to deconvolute its spatial transcriptomics data with matched scRNA-seq data as references to obtain proportions of cell types in the spot. We evaluated the correlation between gene set enrichment scores and proportions of cancer cells.

### Deconvolution of bulk RNA-seq data

We utilized the R package “BayesPrism” [29] to deconvolute the bulk RNA-seq data. According to the developer’s vignette, we first performed data preprocessing. To be specific, the counts matrix and cell type metadata of the PDAC scRNA-seq dataset PRJCA001063 were used as a reference to deconvolute bulk RNA-seq gene expression data. For the PACA-AU and CPTAC-PDAC bulk gene expression data, we first removed their log transformation. Next, we extracted matrixes of posterior means of cell type fractions θ and posterior means of cell type-specific gene expression counts. For downstream analysis, we used the “vst” function from the “DESeq2” R package to normalize the deconvoluted gene expression counts.

### Clustering analysis of bulk RNA-seq data

We identified subgroups of tumor samples in deconvoluted bulk RNA-seq data based on the enrichment scores of CAFs-specific pathways by using the hierarchical clustering method. Before clustering, we normalized the pathway enrichment scores by z-score and transformed them into distance matrices by the R function “dist” with the “method” parameter = “Euclidean.” We implemented the hierarchical clustering with the R function “hclust” in the R package “stats” with the parameters: method = “ward.D2” and members = NULL.

### Evaluation of tumor purity and stromal content

We used the ESTIMATE algorithm [30] to evaluate tumor purity and stromal content (stromal score) for each bulk RNA-seq tumor sample based on its gene expression profile without deconvolution.

### Quantification of copy number alteration (CNA), homologous recombination deficiency (HRD), and intratumor heterogeneity (ITH)

CNA was defined as the number of copy number altered segments in the tumor [31]. The HRD scores were calculated by combining loss of heterozygosity, large-scale state transitions and the number of telomeric allelic imbalances [32]. We used two algorithms (DITHER [33] and DEPTH [34]) to evaluate ITH at the DNA and mRNA level, respectively. DITHER calculates a tumor’s ITH score based on both somatic mutation and CNA profiles [33], and DEPTH evaluates ITH based on gene expression profiles [34]. DITHER scores were calculated using the R package “DITHER” with the input of “maf” and “SNP6” files, and DEPTH scores were calculated using the R package “DEPTH” with the input of gene expression matrix.

### Single-sample gene-set enrichment analysis in bulk RNA-seq data

We utilized the single-sample gene-set enrichment analysis (ssGSEA) [35] to calculate the enrichment scores of gene sets representing biological processes or pathways in tumors, based on their expression profiles of the marker or pathway gene sets. The marker or pathway genes are presented in Supplementary Table S1. The R package “GSVA” was employed to perform the ssGSEA algorithm.

### Pathway enrichment analysis in bulk RNA-seq data

We used the software GSEA [36] (version 4.2.3) to identify the KEGG pathways [21] significantly associated with gene sets, based on gene expression profiles. GSEA evaluates the tendency of the genes in a predefined gene set to be distributed in the gene list ranked with phenotypic correlation, thereby assessing their contributions to the phenotype.

### Survival analysis

We compared the survival, including overall survival (OS), disease-specific survival (DSS), disease-free interval (DFI) and progression-free interval (PFI), between different groups of cancer patients. The Kaplan-Meier (K-M) method [37] was used to show the survival time differences and the log-rank test was used to assess the significance of survival time differences. We utilized the R package “survival” to perform survival analysis.

### Statistical analysis

We used the Mann–Whitney *U* test (one-tailed) to compare two classes of non-normally distributed data, including gene set enrichment scores, tumor purity, stromal scores, CNA scores, HRD scores, ITH scores, and proportions of cell types. To compare two classes of normally distributed data, such as gene expressions in tumors, we used two-tailed Student’s *t* tests. The Fisher’s exact test was utilized to analyze contingency tables. To correct for *p* values in multiple tests, we calculated false discovery rate (FDR) using the Benjamini and Hochberg [38] method.

## Results

### scPathClus identifies two subpopulations of CAFs in PDAC

We applied scPathClus to the scRNA-seq dataset PRJCA001063 [18]. This dataset is a single-cell gene expression matrix 𝑋𝑋 (18,008 × 41,986) which contains 41,986 cells from 24 PDAC patients. The cell types in this dataset had been well annotated by the original publication [18]. Within 50 epochs, the MSE loss of autoencoder showed no apparent decrease (Figure 2A). Based on the pathway enrichment matrix 𝑍𝑍 (13,259 × 41,986), the AE generated a latent feature matrix 𝑍𝑍^′^ (64 × 41,986). We performed an unsupervised clustering of the 41,986 cells based on 𝑍𝑍^′^ (Figure 2B). Taking the original cell type annotation as the ground truth (Figure 2C), we found that the pathway enrichment-based clustering eventually grouped the same cell types together with the overall accuracy as high as 0.98 (Figure 2D and Supplementary Table S2). Except acinar cells which were not distinguished, most of the other cell types had clustering sensitivity and specificity greater than 0.9 (Supplementary Table S2). Mapping patient information into the t-SNE graph (Figure 2E), we found that some clusters of tumor cells (annotated as ductal cell type 2) were patient-specific while other clusters of cell types were not patient-specific. Notably, we have used the raw count matrix to calculate the gene-set enrichment scores for each cell, without any data preprocessing steps. It suggests that the pathway enrichment-based clustering method is more robust than gene expression-based clustering methods for single cells.

**Figure 2.**
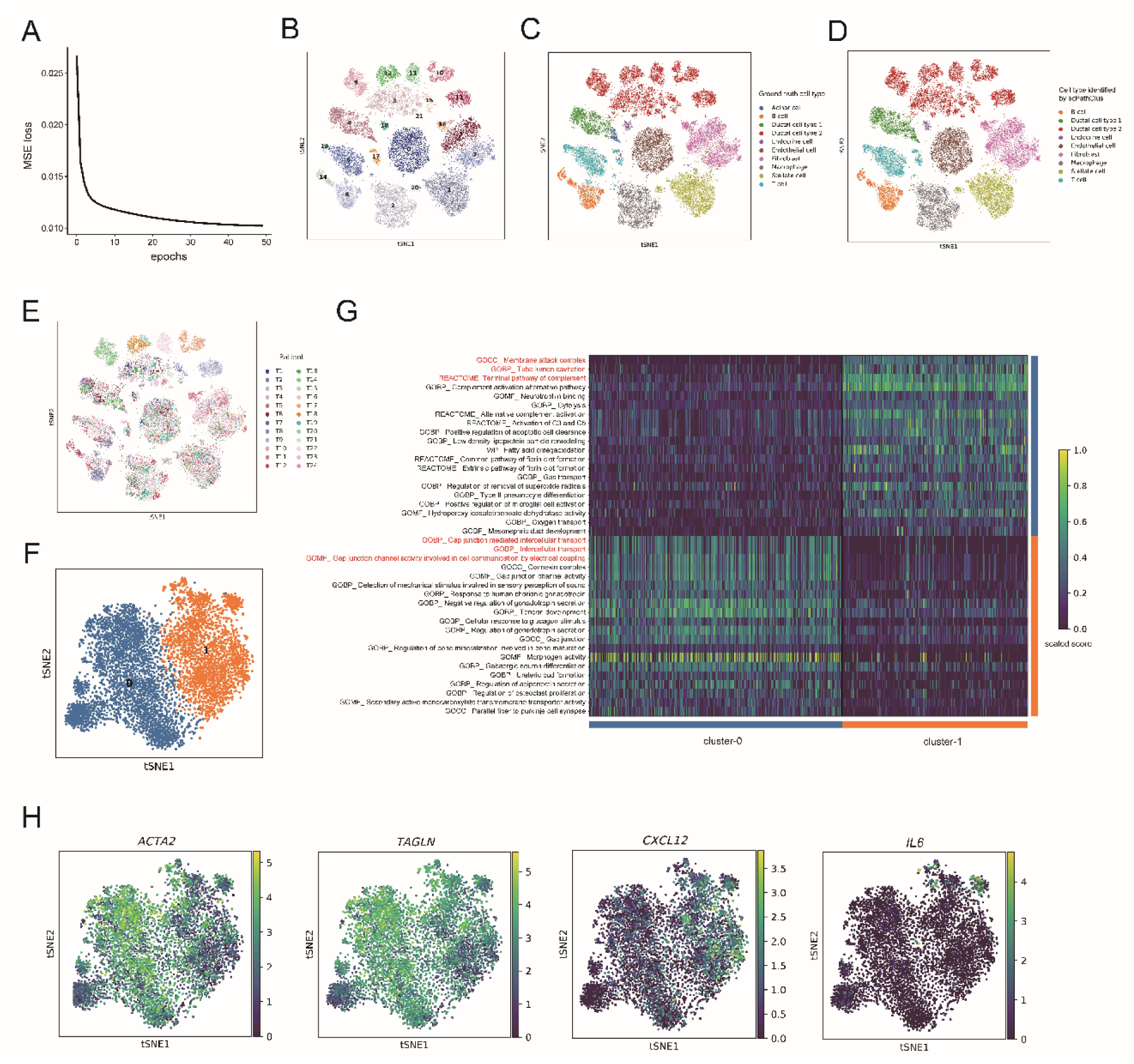
Clustering PDAC single cells by scPathClus identifies two subpopulations of CAFs. **(A)** The mean squared error (MSE) loss within 50 epochs of the autoencoder in the dimension reduction procedure. **(B)** The t-SNE plot showing the 21 cell subclusters identified by scPathClus. **(C)** Mapping the ground truth cell type labels to the t-SNE plot. **(D)** Mapping the cell types obtained by scPathClus to the t-SNE plot. **(E)** Mapping the cell information of patient origin to the t-SNE plot. **(F)** Two subclusters of the CAFs identified by scPathClus through re-clustering of CAFs. **(G)** Heatmap showing the top 20 differentially enriched pathways between the subtypes of CAFs, in which the top 3 marker pathways for each subtype are highlighted in red. The color bar represents the scaled pathway enrichment score. **(H)** t-SNE plots showing the expression level of the marker genes for myCAFs (*ACTA2* and *TAGLN*) and iCAFs (*CXCL12* and *IL6*) in single cells. The color bar represents the normalized gene expression value.

As expected, CAFs showed heterogeneity at the default clustering resolution. Thus, we further clustered CAFs and identified two clusters (resolution = 0.1), termed cluster-0 and cluster-1, respectively (Figure 2F). Furthermore, we performed differential pathway enrichment analysis between cluster-0 and cluster-1 to identify highly active pathways in each cluster using a threshold of FDR < 0.05 and fold change (FC) > 2. This analysis showed that the top 20 pathways with the lowest FDRs highly active in cluster-0 were mainly gap junction-related pathways, while the top 20 pathways highly active in cluster-1 were mainly complement system-related pathways (Figure 2G). Previous studies [9–11] have reported several subtypes of CAFs, such as iCAFs and myCAFs. Chen et al. [39] identified a new CAF subset, namely complement-secreting fibroblasts (csCAFs). Notably, the two CAF subtypes we identified were different from iCAFs and myCAFs (Figure 1H), but incorporated the csCAFs subtype. Here we termed cluster-0 as gapCAFs and cluster-1 as csCAFs, and defined the top 3 highly active pathways in each subtype (with the smallest FDR values) as their specific pathway markers (pathways highlighted in red in Figure 2G). We will discuss the differences between our defined CAF subpopulations and established definition of CAF subpopulations afterwards.

To verify whether the new CAF subtyping method is reproducible, we analyzed another scRNA-seq dataset GSE155698 [19], which contained 48,453 cells from 16 PDAC patients. Likewise, we clustered the 48,453 cells by scPathClus (Supplementary Figure S1A) and annotated their cell types according to the expressions of representative gene markers provided in the original publication [19]. Mapping the t-SNE graph according to the ground truth cell types (Supplementary Figure S1B) and patients (Supplementary Figure S1C) demonstrated scPathClus to successfully cluster the single cells. This analysis identified two subgroups of CAFs (colored in blue) that were far apart in the t-SNE graph (Supplementary Figure S1D), and the differentially expressed pathways between both groups of CAFs were distinct from those between gapCAFs and csCAFs. Because the marker genes *DLN* and *LUM* showed relatively low expression levels in the subgroup of CAFs close to tumor cells (Supplementary Figure S1E), we further analyzed another CAF subgroup that was far from tumor cells by scPathClus. Notably, these CAFs were divided into two major subclusters (Supplementary Figure S1F), which were highly enriched in gap junction-related pathways and complement system-related pathways, respectively (Supplementary Figure S1G).

Altogether, these results support the feasibility and rationality of clustering single cells based on pathway enrichment scores. Furthermore, compared with gene expression-based clustering methods, pathway enrichment-based clustering can detect cell subpopulations with significantly different pathway activity in a more straightforward manner. Using the pathway enrichment-based clustering method, for the first time, we found two types of CAFs showing pathway heterogeneity, as characterized by high activation of the complement system and the gap junction, respectively.

### Both subtypes of CAFs have distinct cell-cell communications with tumor cells in PDAC

We estimated the cell-cell interaction in PRJCA001063 using CellChat [26]. This analysis revealed that both CAF subtypes had ligand-receptor interactions with tumor cells (Figure 3A). Figure 3B showed all significant ligand-receptor interactions between CAFs and tumor cells. Notably, both subtypes of CAFs highly expressed the ligand midkine (MDK), which plays important roles in a range of inflammatory diseases and cancer [40]. Besides, the upregulated ligands secreted by gapCAFs and their paired receptors in tumor cells included MIF - CD74/CD44, SPP1 - CD44, and WNT - FZD5/LRP5. MIF (macrophage migration inhibitory factor) can bind to its homologous receptor (a CD74/CD44 complex) to activate multiple signal transduction pathways, such as Src, ERK, MAPK, and Akt, and to inhibit p53-induced apoptosis [41]; SPP1 interacts with CD44, a cell-surface glycoprotein involved in cell-cell interactions, cell adhesion and migration [42]; WNT interacts with its receptor FZD5 and LRP5 to activate the Wnt signaling pathway [43]. In contrast, the upregulated ligands secreted by csCAFs and their paired receptors in tumor cells included ANGPTL4 - SDC1/SDC4, PTN - SDC1/SDC4/NCL, and NAMPT - INSR. Among them, ANGPTL4 is a Wnt signaling antagonist [44]; PTN is involved in the regulation of cell proliferation, growth and differentiation by acting through various receptors [45]. Taken together, these results suggest distinct crosstalks between both subtypes of CAFs and tumor cells that may lead to different developmental trajectories of tumor cells.

**Figure 3.**
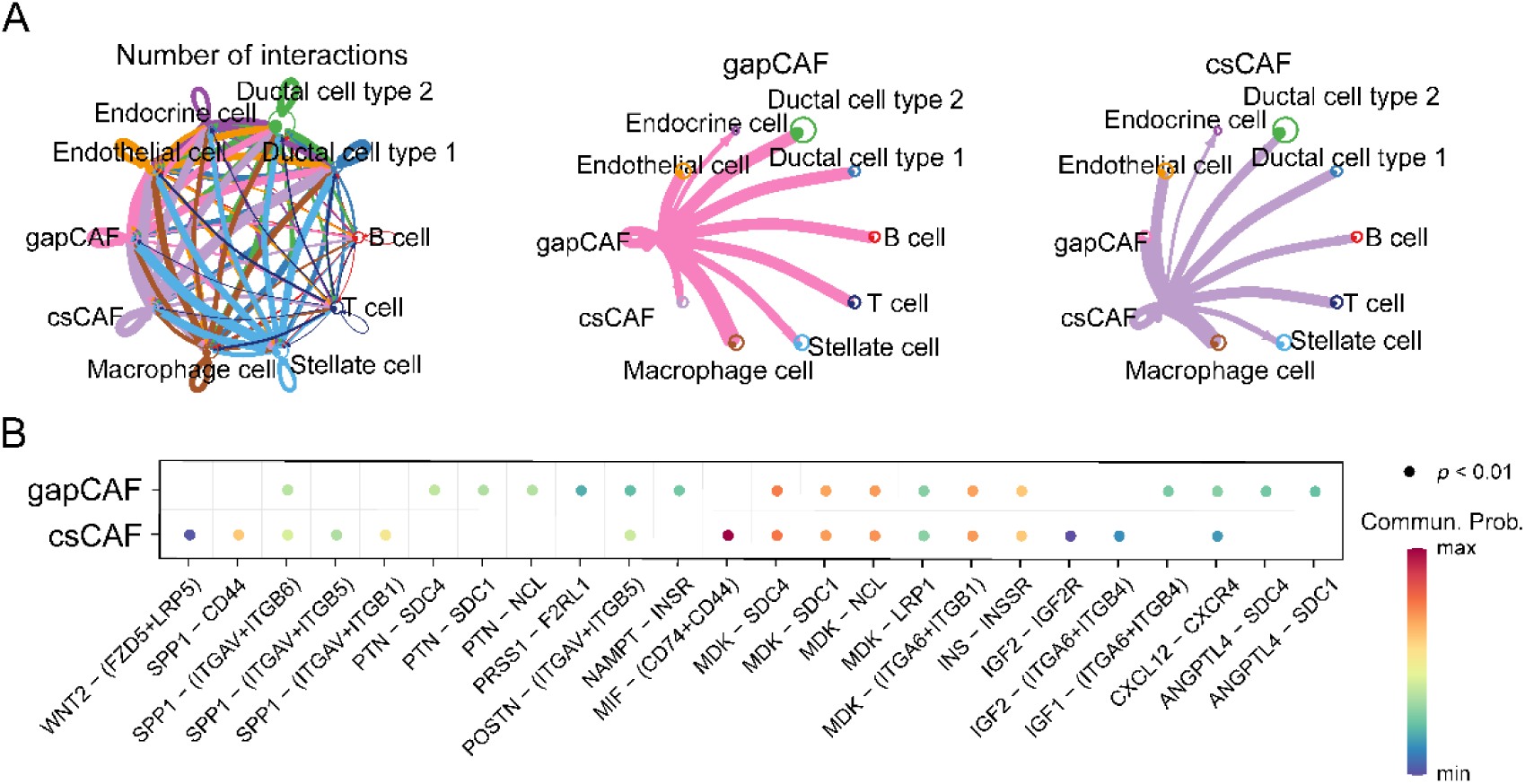
Cell-cell communications between the CAF subpopulations and tumor cells. **(A)** Circle plots displaying the number of ligand-receptor interactions among all cell types in PRJCA001063 PDAC dataset (left), from gapCAFs to tumor cells (ductal cell type 2) (middle) and from csCAFs to tumor cells (right). The number of ligand-receptor interactions is proportional to the thickness of the arrows. **(B)** Bubble plot showing the contribution of significant ligand-receptor pairs between both subtypes of CAFs and tumor cells. The color bar represents the communication probability of ligand-receptor interactions.

### Both subtypes of CAFs display distinct evolutionary trajectories in PDAC

We inferred the evolutional trajectories of both CAF subpopulations in PRJCA001063 using Monocle2 [27]. This analysis revealed that csCAFs occupied an earlier pseudotime branch, while gapCAFs showed multiple cell states later in pseudotime branches (Figure 4A-4C). It implies a relative progression from csCAFs towards gapCAFs. Next, we mapped the enrichment scores of gapCAFs and csCAFs-specific pathways to the pseudotime trajectory. As expected, cells at later pseudotime branches corresponded to higher gapCAFs-specific pathway enrichment scores (Figure 4D), while cells at earlier pseudotime branches corresponded to higher csCAFs-specific pathway enrichment scores (Figure 4E). These results collectively suggest a potential differentiation trajectory from csCAFs to gapCAFs, with gapCAFs being more terminally differentiated and more plastic versus csCAFs.

**Figure 4.**
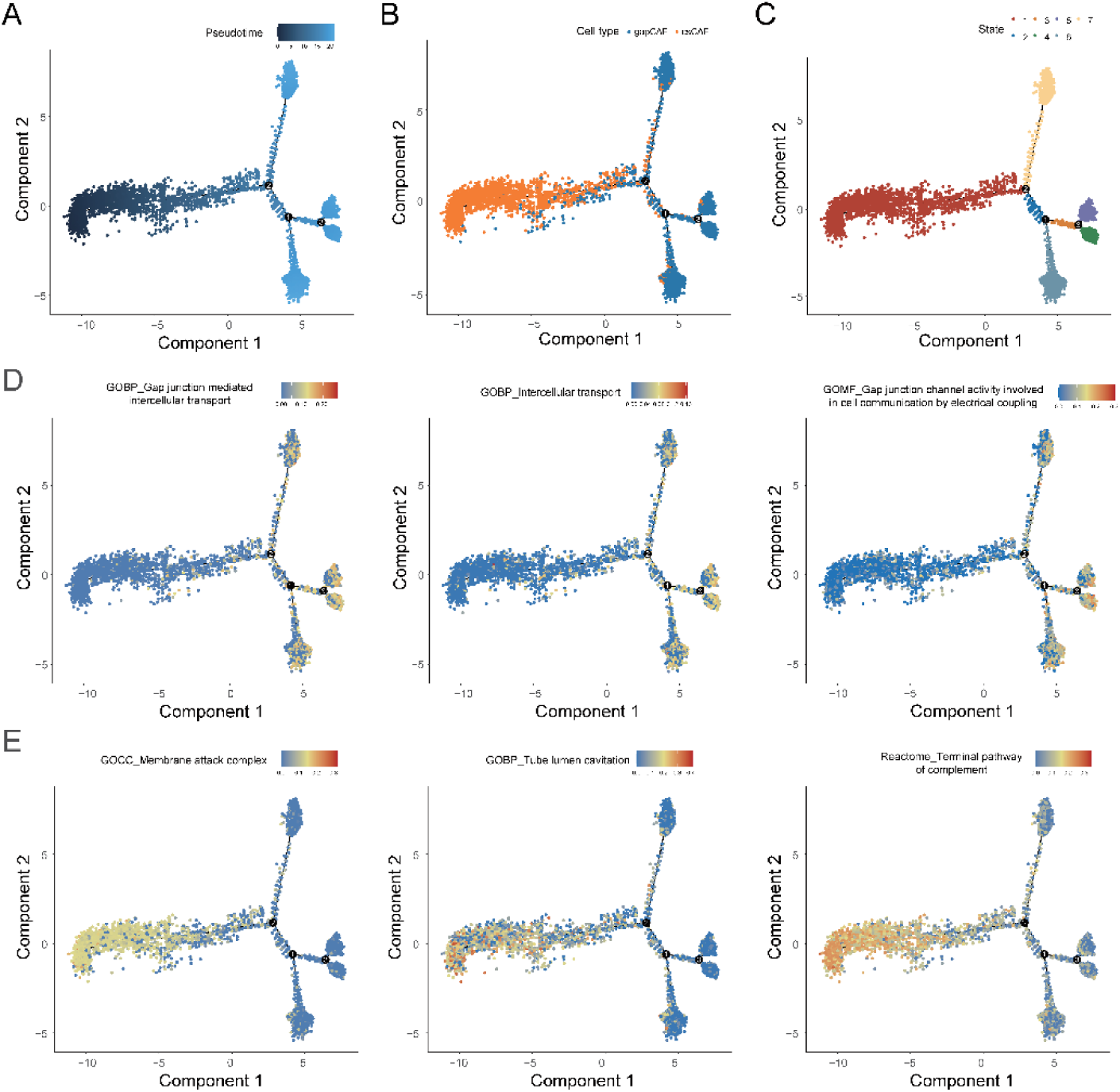
Inferred evolutional trajectories of both CAF subpopulations. **(A)** Pseudotime analysis for both CAF subpopulations in the PRJCA001063 PDAC dataset. **(B)** Regeneration time point overlayed on the pseudotime trajectory for gapCAFs and csCAFs. **(C)** Cell states inferred by Monocle2. **(D)** Mapping the enrichment scores of gapCAF marker pathways in the pseudotime trajectory. **(E)** Mapping the enrichment scores of csCAF marker pathways in the pseudotime trajectory.

### Both subtypes of CAFs display different spatial distribution in PDAC

Previous studies have shown that spatial localization of tumor microenvironment components may result in tumor heterogeneity and offer prognostic value [46]. To observe whether both subtypes of CAFs have different spatial distribution in the tumor, we analyzed a PDAC spatial transcriptome dataset with reference to a paired scRNA-seq dataset [20]. We first deconvolved the spatial transcriptome data using the CARD algorithm [28] to get the cell type composition of each spot. This analysis captured the isolation between cancer (colored in orange) and non-cancer regions, and fibroblasts (colored in blue) were detected in both regions (Figure 5A). We then used the UCell algorithm [24] to calculate the gapCAFs and csCAFs-specific pathway enrichment scores in each spot (Figure 5B). The results showed that the gapCAFs marker pathways were highly activated in the cancer region, while the csCAFs marker pathways were highly activated in the non-cancer region. Spearman correlation analysis showed that the enrichment scores of the gapCAFs marker pathways correlated positively with the proportions of tumor cells in the spots (*p* < 0.001), while csCAFs showed an inverse correlation (*p* < 0.001) (Figure 5C). Taken together, these results suggest that gapCAFs and csCAFs have different spatial distribution in PDAC, with gapCAFs and csCAFs likely located at cancer and non-cancer regions, respectively.

**Figure 5.**
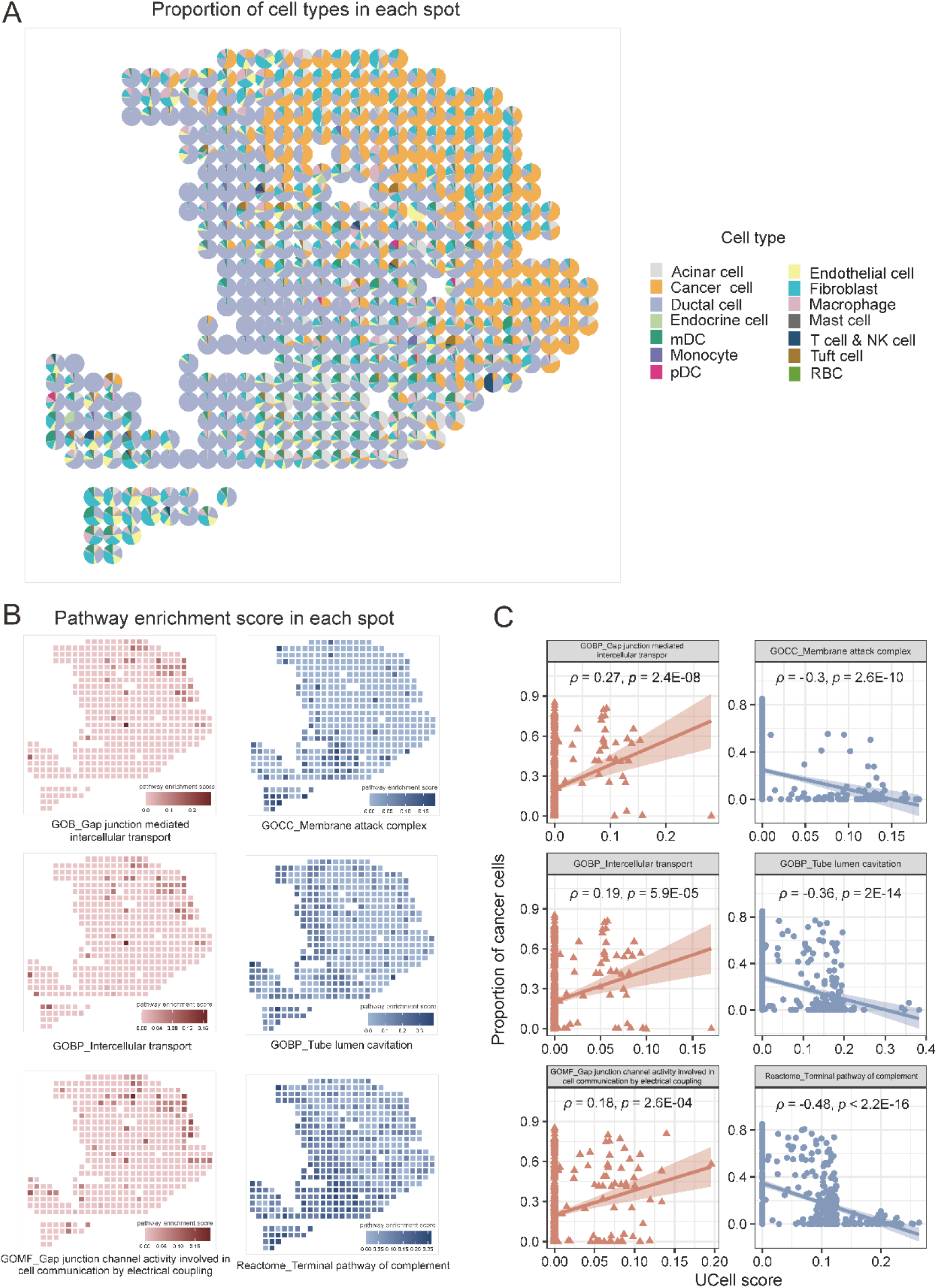
Spatial localization of the CAF subpopulations in PDAC by spatial transcriptome analysis. **(A)** The scatterpie plot displaying the proportions of various cell types in each spot of the spatial transcriptome data. **(B)** Mapping the enrichment scores of gapCAFs marker pathways (left) and csCAFs marker pathways (right) in each spot of the spatial transcriptome data. **(C)** The enrichment scores of gapCAFs and csCAFs marker pathways showing significant positive and negative correlations with the proportions of tumor cells in the spots, respectively. The Spearman correlation coefficients and *p* values are shown.

### Both subtypes of CAFs are associated with different clinical outcomes in PDAC

Although single-cell data have advantages in exploring the heterogeneity of tumors and their microenvironment, most single-cell datasets incorporate small cohorts that does not favor the investigation of the association between cell subpopulations and phenotypic and clinical features. However, bulk transcriptomes often involve large cohorts to fill this gap. Here we used the BayesPrism algorithm [29] to infer the composition of cell types and cell type-specific gene expression profiles in bulk RNA-seq data with the reference to scRNA-seq data.

In the TCGA-PDAC cohort, deconvolution analysis showed that each of the 146 tumor samples contained CAFs, and the proportion of CAFs was greater than 20% in 81 samples (Figure 6A). Next, based on the CAFs-specific gene expression profiles and the enrichment scores of three gapCAFs and three csCAFs marker pathways, we clustered the TCGA-PDAC cohort. This analysis identified three clusters of tumor samples: cluster-1, cluster-2, and cluster-3 (Figure 6B). Notably, the csCAFs and gapCAFs marker pathways were overrepresented in cluster-1 and cluster-3, respectively, indicating that cluster-1 and cluster-3 involved csCAFs-enriched and gapCAFs-enriched tumors, respectively. Survival analysis showed that cluster-1 had significantly better survival prognosis than cluster-3 (*p* = 0.003, 0.008, 0.008 and 0.0003 for OS, DSS, DFI and PFI, respectively) (Figure 6C). Moreover, cluster-3 contained a significantly higher proportion of high-grade (G3&G4) tumors than cluster-1 (54% versus 25%; Fisher’s exact test, *p* = 0.011) (Figure 6D). These results collectively suggest that csCAFs-enriched tumors have more favorable clinical outcomes than gapCAFs-enriched tumors in PDAC.

**Figure 6.**
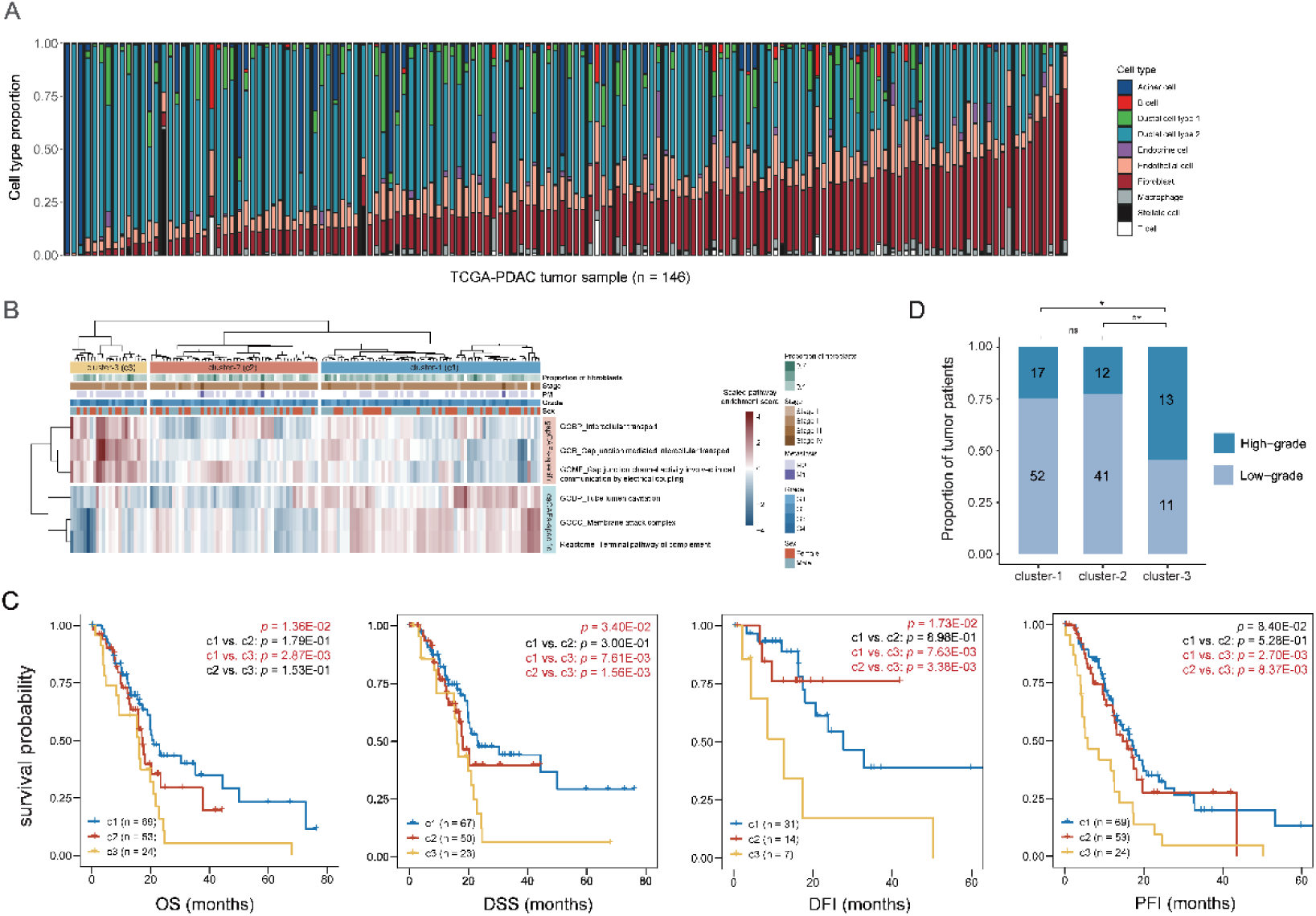
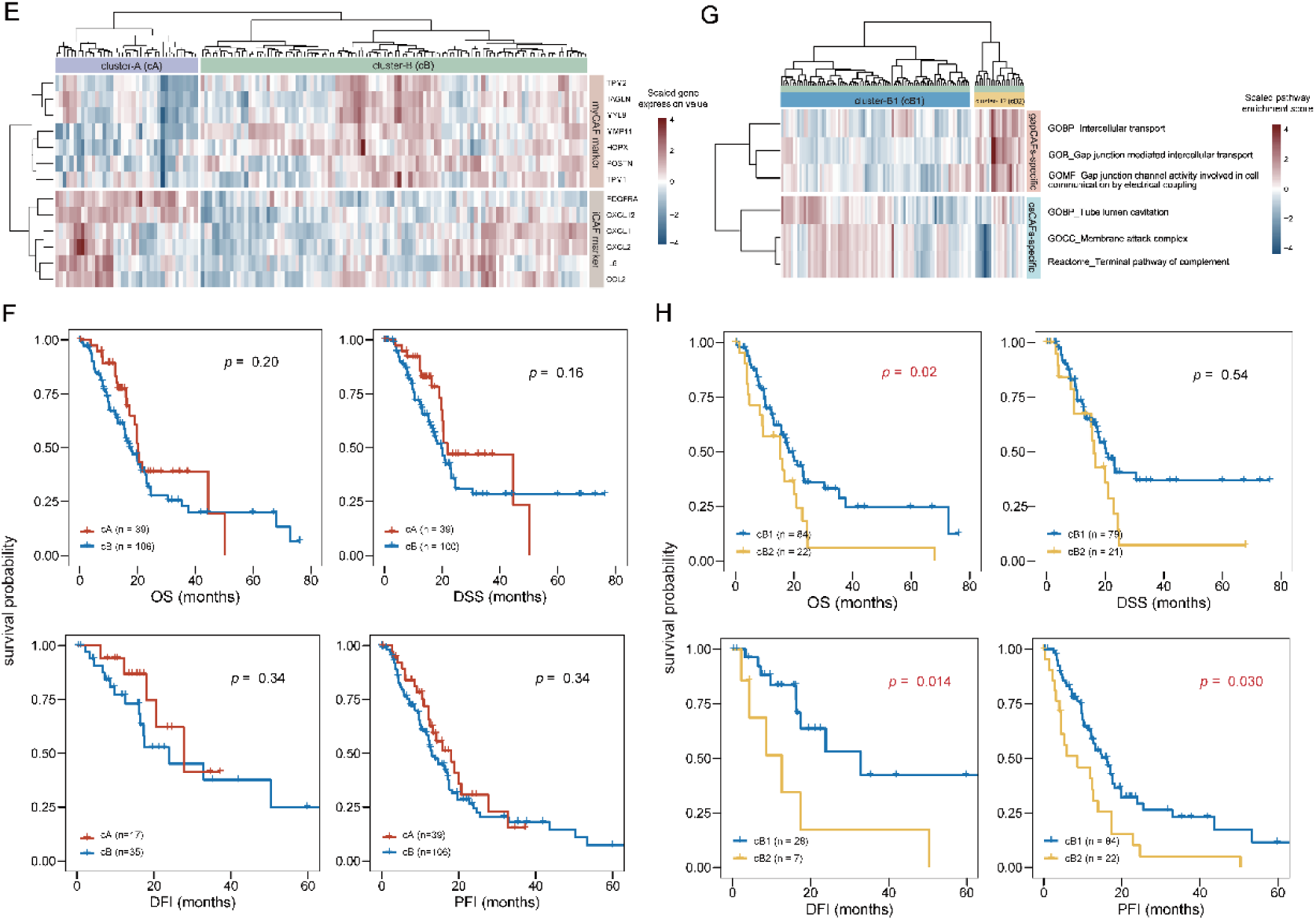
Association between both subtypes of CAFs and clinical outcomes in PDAC. **(A)** The stack bar plot showing the proportions of various cell types obtained by deconvolution analysis of the TCGA-PDAC bulk RNA-seq dataset using the BayesPrism algorithm. The tumor samples were ordered according to the proportion of their fibroblasts. **(B)** Heatmap showing the three PDAC tumor clusters identified by hierarchical clustering using the deconvoluted fibroblast expression profiles based on the enrichment scores of three gapCAFs marker pathways and three csCAFs marker pathways. **(C)** K-M curves showing that patients in cluster-3 have the worst survival while patients in cluster-1 have the best survival. OS, overall survival. DSS, disease-specific survival. DFI, disease-free interval. PFI, progression-free interval. The log-rank test *p* values are shown. **(D)** The stack bar plot showing that cluster-3 contains a significantly higher proportion of high-grade (G3&G4) patients than cluster-1. Fisher’s exact test *p* values are shown. * *p* < 0.05, ** *p* < 0.01, *** *p* < 0.001, ^ns^ not significant; they also apply to the following figures. **(E)** Heatmap showing two TCGA PDAC tumor clusters identified by hierarchical clustering using the deconvoluted fibroblast expression profiles based on the expressions of myCAF and iCAF marker genes. **(F)** K-M curves showing that there is no significant survival difference between both subclusters of PDAC patients identified based on the expressions of myCAF and iCAF marker genes. **(G)** Re-clustering of the myCAFs-enriched tumors identifies csCAFs-enriched and gapCAFs-enriched subclusters. **(H)** K-M curves show significant survival differences between the patients with csCAFs-enriched tumors and the patients with gapCAFs-enriched tumors within the myCAFs-enriched tumors.

As aforementioned, myCAFs and iCAFs were established CAF subtypes [10]. Based on the expressions of their marker genes, we could indeed detect the myCAFs and iCAFs subtypes in the TCGA tumors (Figure 6E and Supplementary Figure S2A). However, both subtypes of tumors showed no significant difference in survival prognosis (Figure 6F and Supplementary Figure S2B). Next, we tried to stratify myCAFs and iCAFs tumors using the marker pathways of gapCAFs and csCAFs. Interestingly, we detected gapCAFs and csCAFs subtypes within myCAFs tumors (Figure 6G), in which csCAFs-enriched tumors still showed significantly better survival than gapCAFs-enriched tumors (Figure 6H). These results suggest that our CAF subtyping method is superior to the established method in light of prognostic value.

We further explored the association of the CAF subtypes with survival prognosis in two other PDAC bulk datasets: CPTAC-PDAC and ICGC-PACA-AU. Likewise, we first performed deconvolution analysis of the bulk transcriptomes and then clustering analysis of the deconvoluted data (Supplementary Figure S2C). Consistent with the prior results, the cluster highly expressing the csCAFs marker pathways showed significantly better OS prognosis than the cluster highly expressing the gapCAFs marker pathways (*p* < 0.05) (Supplementary Figure S2D and Figure S2E). Again, the myCAFs and iCAFs-enriched subtypes in both datasets showed no significant survival difference, while stratifying myCAFs or iCAFs tumors using the marker pathways of gapCAFs and csCAFs detected the subclusters with significant survival difference (Supplementary Figure S2F).

Taken together, these data suggest that the CAF subtypes we identified have significant prognostic value that is absent for the canonic subtypes of CAFs.

### Both subtypes of CAFs are associated with significantly different tumor microenvironment in PDAC

We obtained the proportions of various cell types by deconvolution analysis using the BayesPrism algorithm [29] in the TCGA-PDAC cohort, and compared them among the three clusters identified based on the enrichment scores of gapCAFs and csCAFs marker pathways (Figure 7A). Although the overall proportion of immune cells was low in PDAC, the proportions of macrophages, T cells, and B cells were significantly higher in cluster-1 than in cluster-3 (*p* < 0.01); the proportions of some other cell types like ductal cell type 1 (defined as normal ductal cells), endocrine cells and endothelial cells were also significantly higher in cluster-1 (*p* < 0.05). However, the proportion of tumor cells was significantly lower in cluster-1 than in cluster-3 (*p* = 0.0003). Notably, the proportion of CAFs did not differ significantly among the three clusters. Next, we used the ESTIMATE algorithm [30] to calculate stromal score, immune score, and tumor purity for each tumor sample based on the original gene expression profiles in TCGA-PDAC. We found cluster-1 having significantly higher stromal and immune scores but lower tumor purity than cluster-2 and cluster-3 (*p* < 0.01) (Figure 7B). This result supports the previous finding that gapCAFs are more located at the cancer region and csCAFs more located at non-cancer regions, as cluster-1 and cluster-3 are highly enriched in the csCAFs and gapCAFs marker pathways, respectively. This result also implies why csCAFs-enriched tumors have better prognosis than gapCAFs-enriched tumors since the former confronts a stronger antitumor immune response than the latter.

**Figure 7.**
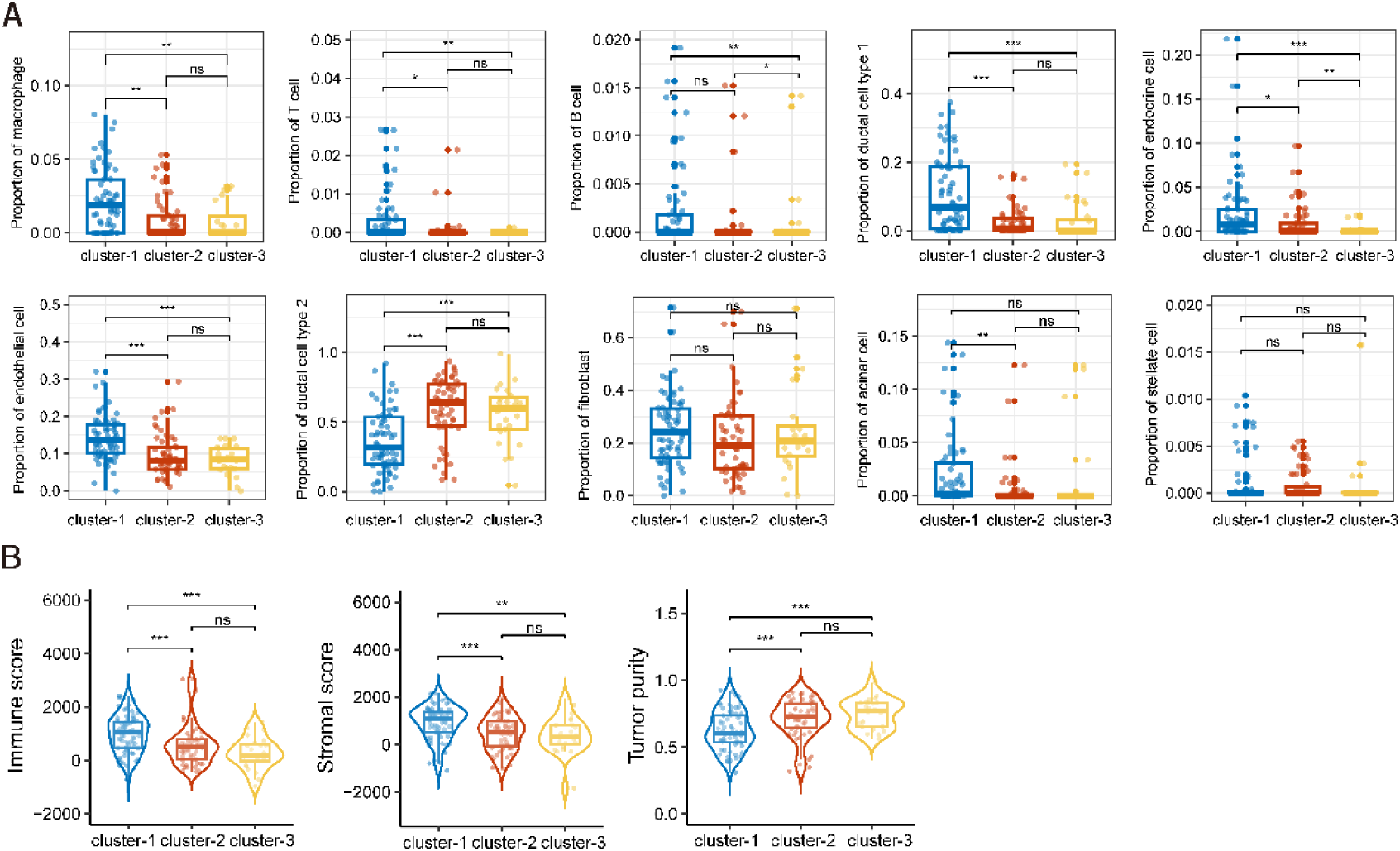
Both subtypes of CAFs are associated with significantly different tumor microenvironment in PDAC. **(A)** Comparisons of the proportions of various cell types among the three clusters identified by pathway enrichment-based clustering of the TCGA-PDAC tumors. **(B)** The csCAFs-enriched tumors (cluster-1) have the highest stromal and immune scores but lowest tumor purity among the three TCGA PDAC clusters. The one-tailed Mann– Whitney *U* test *p* values are indicated in (**A, B**).

### Both subtypes of CAFs are associated with significantly different molecular and phenotypic features in PDAC

We compared tumor-associated molecular and phenotypic features between csCAFs- enriched and gapCAFs-enriched tumors, including genomic instability, ITH, proliferation, stemness and cancer-associated signaling. Genomic instability plays a key role in tumor initiation and progression [47]. We compared the representative features of genomic instability among the three clusters identified based on the marker pathways of gapCAFs and csCAFs in the TCGA-PDAC cohort. HRD may lead to large-scale genomic instability [32]. We found that HRD scores were significantly higher in cluster-3 than in cluster-1 and cluster-2 (*p* < 0.05) (Figure 8A). Tumor aneuploidy, also known as CNA, often results from genomic instability [48], which showed significantly higher levels in cluster-3 than in cluster-1 (*p* = 0.037) (Figure 8B). DNA repair pathways are crucial in maintaining genomic instability [49]. Consistently, the enrichment scores of six key DNA repair pathways (mismatch repair, base excision repair, nucleotide excision repair, Fanconi anemia, homology-dependent recombination, and direct repair) were significantly higher in cluster-3 than in cluster-1 (*p* < 0.05) (Figure 8C). These results collectively suggest that gapCAFs-enriched tumors are more genomically instable than csCAFs-enriched tumors in PDAC.

**Figure 8.**
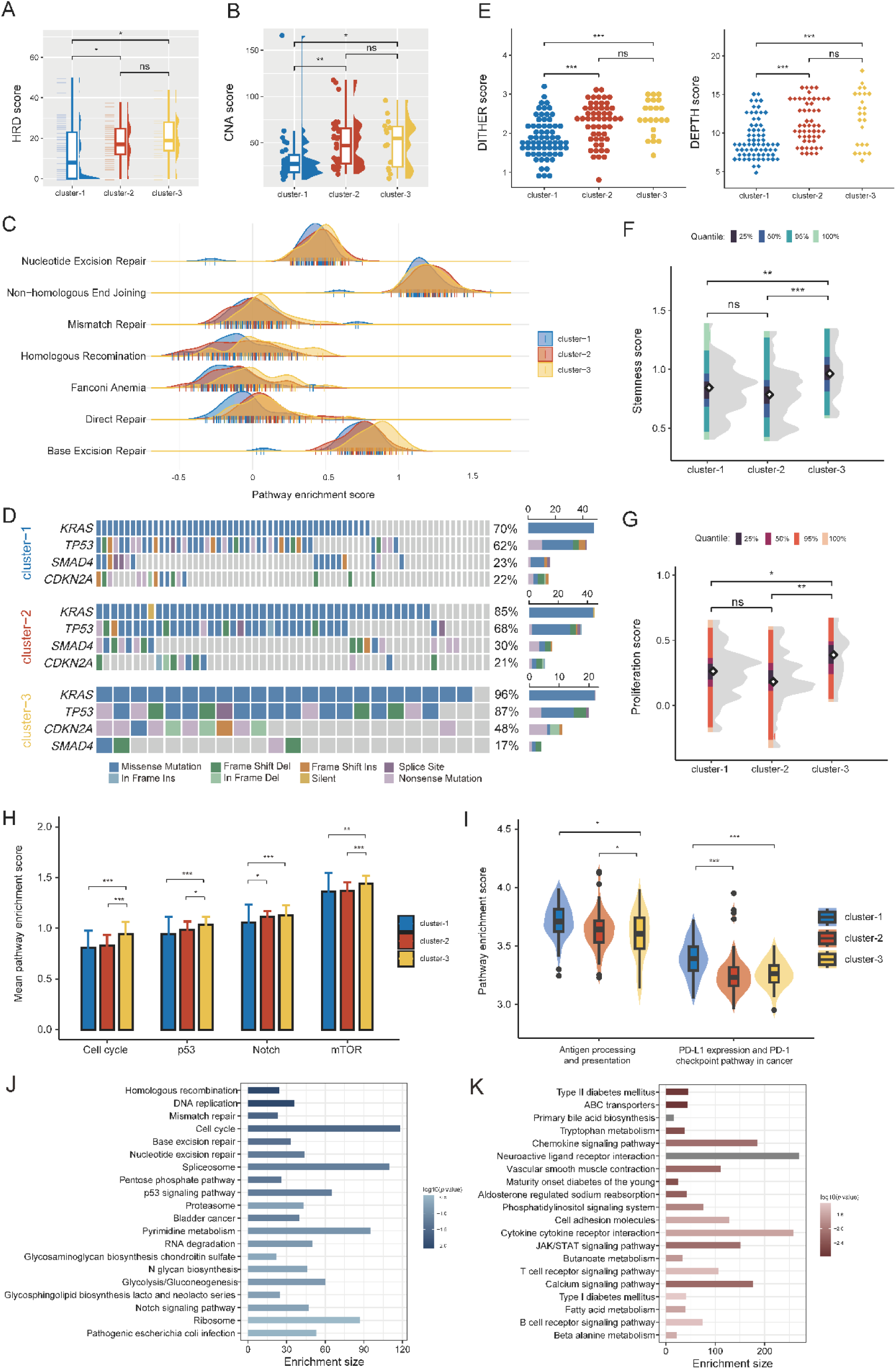
Both subtypes of CAFs display different molecular and phenotypic features in PDAC. **(A, B)** The csCAFs-enriched tumors (cluster-1) have the lowest HRD scores **(A)** and CNA scores **(B)** among the three TCGA PDAC clusters. **(C)** The enrichment scores of six DNA damage repair pathways are significantly higher in gapCAFs-enriched tumors (cluster-3) than in csCAFs-enriched tumors (cluster-1). **(D)** Mutation frequencies of four key genes in the three TCGA PDAC clusters. **(E)** ITH scores (DITHER scores and DEPTH scores) are the lowest in csCAFs-enriched tumors (cluster-1). **(F, G)** gapCAFs-enriched tumors (cluster-3) have the highest stemness scores **(F)** and proliferation scores **(G)**. **(H, I)** gapCAFs-enriched tumors (cluster-3) have significantly higher oncogenic pathway enrichment scores **(H)** but lower immune-related pathway enrichment scores than csCAFs-enriched tumors (cluster-1) **(I)**. **(J, K)** The top 20 KEGG pathways with the highest contribution to the cluster-3 phenotype **(J)** and the cluster-1 phenotype **(K)**, respectively.

The driving mutations of four key genes (*KRAS*, *TP53*, *CDKN2A* and *SMAD4*) play important roles in pancreatic cancer occurrence and progression [50]. Notably, *KRAS*, *TP53* and *CDKN2A* showed significantly higher mutation rates in cluster-3 than in cluster-1 (*KRAS*: 96% versus 70%; *TP53*: 87% versus 62%; *CDKN2A*: 48% versus 22%; Fisher’s exact test, *p* < 0.05) (Figure 8D). Specially, the mutation rate of *KRAS* in cluster-3 was spectacularly as high as 96%. ITH and stemness are two typical properties of cancer that are associated with tumor progression, recurrence, and therapeutic resistance [51, 52]. We found that both DNA and mRNA-level ITH scores (DITHER scores [33] and DEPTH scores [34]) were significantly higher in cluster-3 than in cluster-1 (Figure 8E). Likewise, cluster-3 had significantly higher stemness scores than cluster-1 (*p* = 0.009) (Fig. 8F). Besides, cluster-3 displayed significantly higher proliferation signature scores than cluster-1 (*p* = 0.01) (Fig.8G). Furthermore, we found the enrichment levels of numerous oncogenic pathways, including the cell cycle, p53, Notch, and mTOR signaling pathways, were significantly higher in cluster-3 than in cluster-1 (*p* < 0.01) (Fig. 8H). However, the enrichment levels of immune-related pathways, such as antigen processing and presentation and PD-L1 expression and PD- 1 checkpoint pathway in cancer, were significantly higher in cluster-1 than in cluster-3 (*p* < 0.05) (Fig. 8I).

Taken together, these results demonstrate that gapCAFs-enriched tumors are more endowed with tumor-promoting characteristics and malignant phenotypes compared to csCAFs-enriched tumors. It is in line with the previous findings that: (1) gapCAFs- enriched tumors have worse prognosis than csCAFs-enriched tumors; (2) gapCAFs are largely located at the cancer region and csCAFs are largely located at non-cancer regions; and (3) csCAFs-enriched tumors meet with stronger antitumor immune responses than gapCAFs-enriched tumors.

We further identified the KEGG pathways highly enriched in cluster-1 and cluster-3, respectively, using the GSEA software [36]. The pathways highly enriched in cluster- 3 included cell cycle, DNA repair, glycolysis/gluconeogenesis, spliceosome, RNA degradation, p53 signaling, and Notch signaling (Figure 8J). The pathways highly enriched in cluster-1 included B/T cell receptor signaling, cytokine-cytokine receptor interaction, chemokine signaling, cell adhesion molecules, JAK/STAD signaling, calcium signaling, neuroactive ligand receptor interaction, vascular smooth muscle contraction, fatty acid/butanoate/beta alanine/tryptophan metabolism, and diabetes-related pathways (Figure 8K). Overall, these results confirmed that gapCAFs-enriched tumors highly express cancer-associated pathways, while csCAFs-enriched tumors highly express immune and stromal pathways.

## Discussion

To date, all orthodox immunotherapies have been less effective for PDAC compared to other malignant tumors, and PDAC often shows resistance to cytotoxic chemotherapies with rapid progression [53]. As an important component of the TME, CAFs have been explored as diagnostic and prognostic markers in pancreatic cancer [54–56]. Here we proposed scPathClus, a pathway-based clustering method for single cells. Using scPathClus, we identified two CAF subpopulations (gapCAFs and csCAFs) in PDAC, which show high heterogeneity in pathway activity. Our method overcomes the limitation of existing single-cell clustering and pathway analysis methods, as existing methods cluster single cells based on their gene expressions that are susceptible to dropout events in high-throughput scRNA-seq data. We argue that clustering single cells based on pathway or gene set enrichment scores can alleviate dropout events of single genes’ expression in single cells. Besides, scPathClus uses a gene set scoring method [24] that depends on the relative gene expression in individual cells and thus is robust to data composition; scPathClus uses AE to eliminate noise and redundancy on important features so that the feature selection process is skipped; furthermore, compared to the linear transformation of principal component analysis (PCA), AE can better capture the underlying nonlinear relationship between signaling pathways and cells.

Although there have already been some subtyping methods for CAFs, such as myCAFs expressing marker genes *ACTA2* and *TAGLN* and iCAFs characterized by high expression of *IL6* and *CXCL12.* In this study, we used scPathClus to classify CAFs and identified two subpopulations (gapCAFs and csCAFs) that are different from myCAFs and iCAFs. Interestingly, although gapCAFs had certain overlaps with myCAFs, both types of CAFs display contrastive properties⸺myCAFs play a role in tumor suppression [9–11] while gapCAFs have a tumor-promoting function. Notably, the CAF subtypes we identified have prognostic value, as compared with the established subtypes (myCAFs and iCAFs) showing no prognostic difference. Pathway enrichment analysis showed that gapCAFs have high activity of gap junction-related pathways, while csCAFs have high activity of the complement system. Gap junctions, which are composed of connexins, directly connect cells and provide channels for direct information exchange between cells. Connexins have been characterized as tumor suppressors previously [57, 58], but their pro-tumor functions have been recently reported [59, 60]. The synergistic effect of TME components is key in determining tumor progression, in which connexin channels play a significant role [61]. Metabolism energizes tumor growth, but the resulting hypoxic conditions are destructive to tumor cells. Thus, tumor cells must cope with such a circumstance to survive. It has been reported that fibroblasts are connected by gap junctions formed by proteins like connexin-43, to allow hydrogen ions produced by tumor cells to be transmitted across the stromal syncytium [62]. In spheroids of pancreatic cancer, gap junctions between hypoxic and normoxic cells may regulate the acid-base balance at the hypoxic core [63]. The Warburg effect, indicating tumor cells to utilize glucose by glycolysis, is recognized as a major feature of tumors [64]. Our analysis indicates that the glycolysis/gluconeogenesis pathway is enriched in the tumors with high activity of gapCAFs marker pathways. Moreover, spatial transcriptome analysis shows gpCAFs to be situated in the center of tumor regions. Based on these findings, we contend that gpCAFs play a key role in modulating the acidic environment generated by tumor cells with a high metabolism, thereby promoting tumor cell proliferation. However, the dual role of gap junctions in tumors should be noted. For example, Connexin-43 expression on cancer cells has been reported to increase cell permeability to chemotherapeutics [65], while some studies showed that using connexin-43 inhibitors to block gap junctions could prevent the transfer of essential compounds between cells and thus lead to cellular apoptosis [66, 67]. As for csCAFs, Chen et al. [39] demonstrated them located in the tissue stroma adjacent to malignant ductal cells only in early PDAC, and suggested csCAFs playing a role of PDAC inhibition. Our pathway enrichment-based clustering analysis confirmed csCAFs to be a subtype of CAFs in PDAC. There is evidence that complement proteins are present in the TME, and tumor cells, immune cells and stromal cells are able to produce large amounts of these components in situ [68]. In addition to the function of membrane attacking, csCAFs may regulate the PDAC immune microenvironment, as evidenced by our data. We found that csCAFs- enriched tumors involved a significantly higher ratio of immune cells and highly expressed immune-related pathways, such as the antigen processing and presentation and the PD-L1 expression and PD-1 checkpoint pathway in cancer. This finding suggests that the abundance of csCAFs is a potential biomarker for immunotherapy.

In conclusion, this work proposed a pathway-based single cells clustering method and applied this method to analyze PDAC single cells. This analysis identified two types of CAFs characterized by high complement system activity and high gap junction pathway activity, respectively. By further analyzing cell-cell communications, spatial transcriptomics and bulk transcriptomics, we demonstrated that both types of CAFs are distinct in their associations with tumor molecular, phenotypic, and clinical features, as well as in spatial distribution. Compared to established CAF subtyping methods, our method displays better prognostic relevance. Thus, this work provides novel insights into the role of CAFs in PDAC and have potential clinical implications for the diagnosis, prognosis and treatment of PDAC patients.

## Declarations

### Ethics approval and consent to participate

Not applicable.

### Consent for publication

Not applicable.

### Availability of data and materials

All data associated with this study are available within the paper and its supplementary data. The codes and scripts of the scPathClus algorithms are available at the website: https://github.com/WangX-Lab/scPathClus.

## Acknowledgments

Not applicable.

## Author Contributions

**Hongjing Ai**: Methodology, Software, Validation, Formal analysis, Investigation, Data curation, Visualization, Writing - original draft. **Rongfang Nie**: Methodology, Investigation. **Xiaosheng Wang**: Conceptualization, Methodology, Resources, Investigation, Writing - original draft, Supervision, Project administration.

## List of Abbreviations

PDAC: pancreatic ductal carcinoma
CAFs: tumor-associated fibroblasts
scRNA-seq: single-cell RNA sequencing
TME: tumor microenvironment
myCAFs: myofibroblasts
iCAFs: inflammatory fibroblasts
apCAFs: antigen-presenting fibroblasts
GEO: Gene Expression Omnibus
UCSC: University of California Santa Cruz
ICGC: the International Cancer Genome Consortium
MSigDB: the Molecular Signatures Database
GO: Gene Ontology
AE: autoencoder
MSE: mean squared error
API: application programming interface
CNA: copy number alteration
HRD: homologous recombination deficiency
ssGSEA: single-sample gene-set enrichment analysis
K-M: Kaplan-Meier
OS: overall survival
DSS: disease-specific survival
DFI: disease-free interval
PFI: progression-free interval
FC: fold change
csCAFs: complement-secreting fibroblasts
MDK: midkine
MIF: macrophage migration inhibitory factor
PCA: principal component analysis.

