## Supplemental materials for "Pathway-based clustering identifies two subtypes of cancer-associated fibroblasts associated with distinct molecular and clinical features in pancreatic ductal carcinoma"

Supplementary Materials for

### Pathway enrichment-based unsupervised learning identifies novel subtypes of cancer-associated fibroblasts in pancreatic ductal carcinoma

**This PDF file includes:**

Supplementary Figure S1 to S2
Supplementary Table S1 to S2

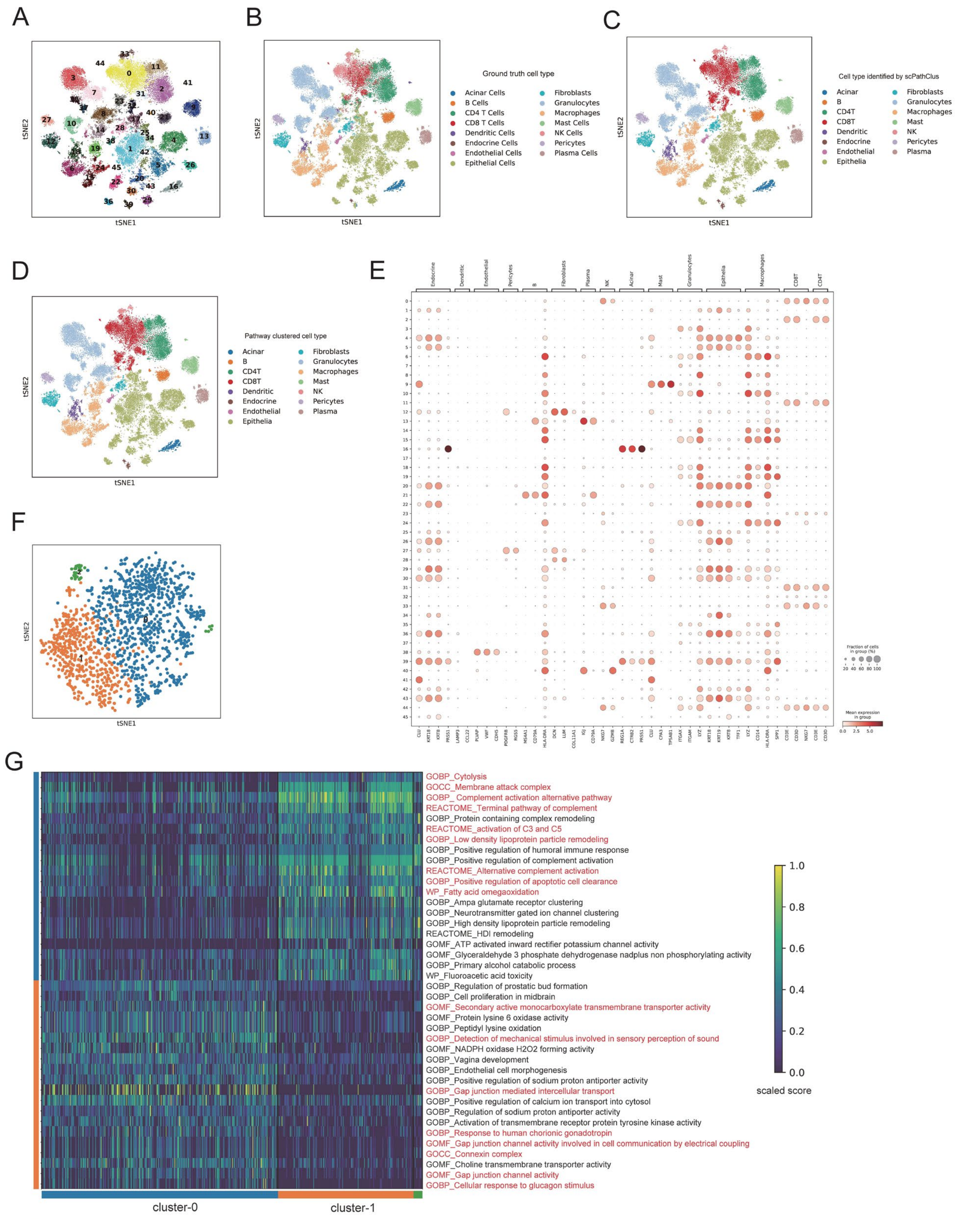
A
B
C
D
E
F
G
tSNE1
tSNE2
Ground truth cell type
Acinar Cells
B Cells
CD4 T Cells
CD8 T Cells
Dendritic Cells
Endocrine Cells
Endothelial Cells
Epithelial Cells
Fibroblasts
Granulocytes
Macrophages
Mast Cells
NK Cells
Pericytes
Plasma Cells
Cell type identified by scPathClus
Pathway clustered cell type
Acinar
B
CD4T
CD8T
Dendritic
Endocrine
Endothelial
Epithelia
Fibroblasts
Granulocytes
Macrophages
Mast
NK
Pericytes
Plasma
GOBP_Cytolysis
GOCC_Membrane attack complex
GOBP_ Complement activation alternative pathway
REACTOME_Terminal pathway of complement
GOBP_Protein containing complex remodeling
REACTOME_activation of C3 and C5
GOBP_Low density lipoprotein particle remodeling
GOBP_Positive regulation of humoral immune response
GOBP_Positive regulation of complement activation
REACTOME_Alternative complement activation
GOBP_Positive regulation of apoptotic cell clearance
WP_Fatty acid omegaoxidation
GOBP_Ampa glutamate receptor clustering
GOBP_Neurotransmitter gated ion channel clustering
GOBP_High density lipoprotein particle remodeling
REACTOME_HDl remodeling
GOMF_ATP activated inward rectifier potassium channel activity
GOMF_Glyceraldehyde 3 phosphate dehydrogenase nadplus non phosphorylating activity
GOBP_Primary alcohol catabolic process
WP_Fluoroacetic acid toxicity
GOBP_Regulation of prostatic bud formation
GOBP_Cell proliferation in midbrain
GOMF_Secondary active monocarboxylate transmembrane transporter activity
GOMF_Protein lysine 6 oxidase activity
GOBP_Peptidyl lysine oxidation
GOBP_Detection of mechanical stimulus involved in sensory perception of sound
GOMF_NADPH oxidase H2O2 forming activity
GOBP_Vagina development
GOBP_Endothelial cell morphogenesis
GOBP_Positive regulation of sodium proton antiporter activity
GOBP_Gap junction mediated intercellular transport
GOBP_Positive regulation of calcium ion transport into cytosol
GOBP_Regulation of sodium proton antiporter activity
GOBP_Activation of transmembrane receptor protein tyrosine kinase activity
GOBP_Response to human chorionic gonadotropin
GOMF_Gap junction channel activity involved in cell communication by electrical coupling
GOCC_Connexin complex
GOMF_Choline transmembrane transporter activity
GOMF_Gap junction channel activity
GOBP_Cellular response to glucagon stimulus
1.0
0.8
0.6
0.4
0.2
0.0
scaled score
cluster-0
cluster-1

**Figure S1.**

**Using scPathClus to cluster the PDAC scRNA-seq data GSE155698 can reproduce the CAF subpopulations.** **(A)** The t-SNE plot showing the subclusters identified by scPathClus. **(B)** Mapping the ground truth cell type labels to the t-SNE plot. **(C)** Mapping the cell types obtain by scPathClus to the t-SNE plot. **(D)** Mapping the cell information of patient-origin to the t-SNE plot. **(E)** Dot plot showing the expression level of cell type marker genes in each subcluster. **(F)** Two subclusters of the CAFs identified by scPathClus through re-clustering. **(G)** Heatmap for the top 20 differentially enriched pathways in the two subtypes of CAFs, and the pathways consistent with those obtained in PDAC_CRA001160 dataset are highlighted in red.

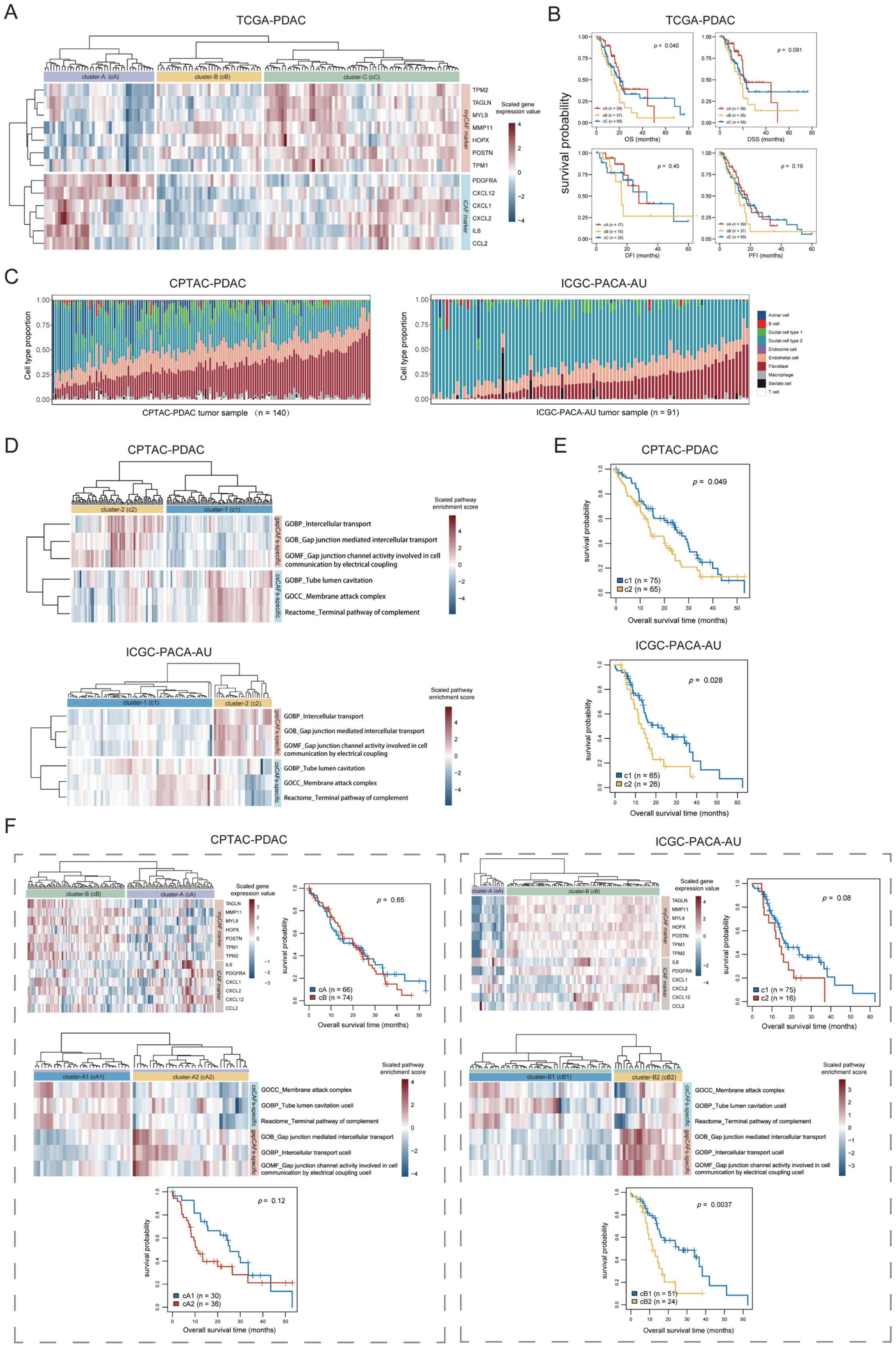
A
TCGA-PDAC
cluster-A (cA)
cluster-B (cB)
cluster-C (cC)
TPM2
TAGLN
MYL9
MMP11
HOPX
POSTN
TPM1
PDGFRA
CXCL12
CXCL1
CXCL2
IL6
CCL2
myCAF marker
iCAF marker
Scaled gene expression value
B
TCGA-PDAC
survival probability
p = 0.040
p = 0.091
p = 0.45
p = 0.19
OS (months)
DSS (months)
DFI (months)
PFI (months)
C
CPTAC-PDAC
ICGC-PACA-AU
Cell type proportion
CPTAC-PDAC tumor sample （n = 140）
ICGC-PACA-AU tumor sample (n = 91)
Acinar cell
B cell
Ductal cell type 1
Ductal cell type 2
Endocrine cell
Endothelial cell
Fibroblast
Macrophage
Stellate cell
T cell
D
CPTAC-PDAC
cluster-2 (c2)
cluster-1 (c1)
GOBP_Intercellular transport
GOB_Gap junction mediated intercellular transport
GOMF_Gap junction channel activity involved in cell communication by electrical coupling
GOBP_Tube lumen cavitation
GOCC_Membrane attack complex
Reactome_Terminal pathway of complement
gapCAFs-specific
csCAFs-specific
Scaled pathway enrichment score
ICGC-PACA-AU
cluster-1 (c1)
cluster-2 (c2)
E
CPTAC-PDAC
p = 0.049
c1 (n = 75)
c2 (n = 65)
survival probability
Overall survival time (months)
ICGC-PACA-AU
p = 0.028
c1 (n = 65)
c2 (n = 26)
F
CPTAC-PDAC
ICGC-PACA-AU
p = 0.65
cA (n = 66)
cB (n = 74)
p = 0.08
c1 (n = 75)
c2 (n = 16)
cluster-A1 (cA1)
cluster-A2 (cA2)
cluster-B1 (cB1)
cluster-B2 (cB2)
GOBP_Tube lumen cavitation ucell
GOBP_Intercellular transport ucell
GOMF_Gap junction channel activity involved in cell communication by electrical coupling ucell
p = 0.12
cA1 (n = 30)
cA2 (n = 38)
p = 0.0037
cB1 (n = 51)
cB2 (n = 24)

**Figure S2.**

**Validation of the relevance between CAF subpopulations and clinical outcomes in PDAC. (A)** Heatmap showing the three clusters of TCGA-PDAC identified by hierarchical clustering based on myCAF and iCAF marker genes. **(B)** K-M curves showing that there is no significant difference in DSS, DFI and PFI between the groups identified by CAFs marker gene. **(C)** The stack bar plots showing the cell type proportions obtained by deconvolution of CPTAC-PDAC (left) and PACA-AU (right) cohorts. **(D)** Heatmap showing the two clusters identified by hierarchical clustering on the deconvoluted fibroblast expression profile based on the enrichment scores of three gapCAF marker pathways and three csCAF marker pathways in CPTAC-PDAC (top) and PACA-AU (bottom) cohorts. **(E)** K-M curves showing that patients with csCAF enriched tumors have the better survival than that with gapCAF enriched tumors in CPTAC-PDAC (top) and PACA-AU (bottom) cohorts. **(F)** PDAC subtypes clustered by myCAF and iCAF marker gene could be stratified again using gapCAF and csCAF marker pathways.

**Supplementary Table S1.**

**The clustering performance of the scPathClus algorithm.**

**PDAC_CRA001160：**

**Overall accuracy**

| Statistical index | Statistical value |
|---|---|
| Accuracy | 0.964 |
| Kappa | 0.957 |
| AccuracyLower | 0.962 |
| AccuracyUpper | 0.965 |
| AccuracyNull | 0.269 |
| AccuracyPValue | 0.000 |
| McnemarPValue | NA |

**Clustering performance for each cell type**

| Class | Sensitivity | Specificity | Pos Pred Value | Neg Pred Value | Precision | Recall |
|---|---|---|---|---|---|---|
| Fibroblast | 0.961 | 0.999 | 0.992 | 0.994 | 0.992 | 0.961 |
| Stellate cell | 0.982 | 0.993 | 0.954 | 0.998 | 0.954 | 0.982 |
| Macrophage | 0.970 | 0.999 | 0.994 | 0.996 | 0.994 | 0.970 |
| Endothelial cell | 0.998 | 0.999 | 0.991 | 1.000 | 0.991 | 0.998 |
| T cell | 0.957 | 0.992 | 0.920 | 0.996 | 0.920 | 0.957 |
| B cell | 0.924 | 0.999 | 0.980 | 0.995 | 0.980 | 0.924 |
| Ductal cell type 2 | 0.993 | 0.996 | 0.989 | 0.997 | 0.989 | 0.993 |
| Endocrine cell | 0.944 | 0.997 | 0.702 | 1.000 | 0.702 | 0.944 |
| Ductal cell type 1 | 0.813 | 0.999 | 0.982 | 0.985 | 0.982 | 0.813 |
| Acinar cell | NA | 0.988 | NA | NA | 0.000 | NA |

| Class | F1 | Prevalence | Detection Rate | Detection Prevalence | Balanced Accuracy |
|---|---|---|---|---|---|
| Fibroblast | 0.976 | 0.143 | 0.137 | 0.138 | 0.980 |
| Stellate cell | 0.968 | 0.122 | 0.120 | 0.126 | 0.988 |
| Macrophage | 0.982 | 0.117 | 0.114 | 0.114 | 0.985 |
| Endothelial cell | 0.994 | 0.121 | 0.121 | 0.122 | 0.998 |
| T cell | 0.938 | 0.083 | 0.079 | 0.086 | 0.975 |
| B cell | 0.951 | 0.061 | 0.056 | 0.058 | 0.962 |
| Ductal cell type 2 | 0.991 | 0.269 | 0.267 | 0.269 | 0.994 |
| Endocrine cell | 0.805 | 0.008 | 0.008 | 0.011 | 0.970 |
| Ductal cell type 1 | 0.890 | 0.076 | 0.062 | 0.063 | 0.906 |
| Acinar cell | NA | 0.000 | 0.000 | 0.012 | NA |

**GSE155698：**

**Overall accuracy**

| Statistical index | Statistical value |
|---|---|
| Accuracy | 0.864 |
| Kappa | 0.838 |
| AccuracyLower | 0.861 |
| AccuracyUpper | 0.867 |
| AccuracyNull | 0.300 |
| AccuracyPValue | 0.000 |
| McnemarPValue | NA |

**Clustering performance for each cell type**

| Class | Sensitivity | Specificity | Pos Pred Value | Neg Pred Value | Precision | Recall | F1 |
|---|---|---|---|---|---|---|---|
| Granulocytes | 0.932 | 0.998 | 0.990 | 0.986 | 0.990 | 0.932 | 0.960 |
| Epithelial | 0.982 | 0.965 | 0.924 | 0.992 | 0.924 | 0.982 | 0.952 |
| Acinar | 0.969 | 0.999 | 0.971 | 0.999 | 0.971 | 0.969 | 0.970 |
| Macrophages | 0.948 | 0.987 | 0.903 | 0.993 | 0.903 | 0.948 | 0.925 |
| NK | 0.579 | 0.948 | 0.093 | 0.996 | 0.093 | 0.579 | 0.160 |
| B | 0.880 | 0.999 | 0.964 | 0.997 | 0.964 | 0.880 | 0.920 |
| CD4 T | 0.939 | 0.975 | 0.784 | 0.994 | 0.784 | 0.939 | 0.854 |
| Mast | 0.943 | 0.999 | 0.977 | 0.998 | 0.977 | 0.943 | 0.960 |
| Plasma | 0.982 | 0.996 | 0.871 | 1.000 | 0.871 | 0.982 | 0.923 |
| CD8 T | 0.376 | 0.995 | 0.912 | 0.916 | 0.912 | 0.376 | 0.533 |
| Dendritic | 0.383 | 0.994 | 0.559 | 0.988 | 0.559 | 0.383 | 0.455 |
| Pericytes | 0.967 | 0.999 | 0.954 | 0.999 | 0.954 | 0.967 | 0.960 |
| Fibroblasts | 0.933 | 0.995 | 0.882 | 0.997 | 0.882 | 0.933 | 0.907 |
| Endothelial | 0.986 | 0.998 | 0.786 | 1.000 | 0.786 | 0.986 | 0.875 |
| Endocrine | 0.357 | 1.000 | 0.893 | 0.997 | 0.893 | 0.357 | 0.510 |

| Class | Prevalence | Detection Rate | Detection Prevalence | Balanced Accuracy |
|---|---|---|---|---|
| Granulocytes | 0.172 | 0.161 | 0.162 | 0.965 |
| Epithelial | 0.300 | 0.295 | 0.319 | 0.973 |
| Acinar | 0.021 | 0.020 | 0.021 | 0.984 |
| Macrophages | 0.114 | 0.108 | 0.119 | 0.967 |
| NK | 0.009 | 0.005 | 0.057 | 0.764 |
| B | 0.021 | 0.019 | 0.019 | 0.940 |
| CD4 T | 0.088 | 0.083 | 0.105 | 0.957 |
| Mast | 0.034 | 0.032 | 0.033 | 0.971 |
| Plasma | 0.026 | 0.025 | 0.029 | 0.989 |
| CD8 T | 0.127 | 0.048 | 0.052 | 0.685 |
| Dendritic | 0.020 | 0.008 | 0.014 | 0.689 |
| Pericytes | 0.015 | 0.015 | 0.015 | 0.983 |
| Fibroblasts | 0.041 | 0.038 | 0.044 | 0.964 |
| Endothelial | 0.006 | 0.006 | 0.008 | 0.992 |
| Endocrine | 0.005 | 0.002 | 0.002 | 0.678 |

**Supplementary Table S2. The marker or pathway genes used in this study.**

| biological signatures/ pathways | Gene set |
|---|---|
| Stemness | *DNMT3B, PFAS, XRCC5, HAUS6, TET1, IGF2BP1, PLAA, TEX10, MSH6, DLGAP5, SKIV2L2, SOHLH2, RRAS2, PAICS, CPSF3, LIN28B, IPO5, BMPR1A, ZNF788, ASCC3, FANCB, HMGA2, TRIM24, ORC1, HDAC2, HESX1, INHBE, MIS18A, DCUN1D5, MRPL3, CENPH, MYCN, HAUS1, GDF3, TBCE, RIOK2, BCKDHB,, RAD1, NREP, ADH5, PLRG1, ROR1, RAB3B, DIAPH3, GNL2, FGF2, NMNAT2, KIF20A, CENPI, DDX1, XXYLT1, GPR176, BBS9, C14orf166, BOD1, CDC123, SNRPD3, FAM118B, DPH3, EIF2B3, RPF2, APLP1, DACT1, PDHB, C14orf119, DTD1, SAMM50, CCL26, MED20, UTP6, RARS2, ARMCX2, RARS, MTHFD2, DHX15, HTR7, MTHFD1L, ARMC9, XPOT, IARS, HDX, ACTRT3, ERCC2, TBC1D16, GARS, KIF7, UBE2K, SLC25A3, ICMT, UGGT2, ATP11C, SLC24A1, EIF2AK4, GPX8, ALX1, OSTC, TRPC4, HAS2, FZD2, TRNT1, MMADHC, SNX8, CDH6, HAT1, SEC11A, DIMT1, TM2D2, FST, GBE1* |
| Proliferation | *CCNB1, CDC20, CDKN3, CDK1, MAD2L1, PRC1, RRM2* |
| Base excision repair pathway | *PARP1, POLB, APEX1, APEX2, FEN1, TDG, TDP1, UNG* |
| Nucleotide excision repair pathway | *CUL5, ERCC1, ERCC2, ERCC4, ERCC5, ERCC6, POLE, POLE3, XPA, XPC* |
| Mismatch repair pathway | *EXO1, MLH1, MLH3, MSH2, MSH3, MSH6, PMS1, PMS2* |
| the Fanconi anemia pathway | *FANCA, FANCB, FANCC, FANCD2, FANCI, FANCL, FANCM, UBE2T* |
| Homologous recombination pathway | *MRE11A, NBN, RAD50, TP53BP1, XRCC2, XRCC3, BARD1, BLM, BRCA1, BRCA2, BRIP1, EME1, GEN1, MUS81, PALB2, RAD51, RAD52, RBBP8, SHFM1, SLX1A, TOP3A* |
| Non-homologous end joining pathway | *LIG4, NHEJ1, POLL, POLM, PRKDC, XRCC4, XRCC5, XRCC6* |
| Direct repair pathway | *ALKBH2, ALKBH3, MGMT* |
| Cell cycle | *ABL1, ANAPC1, ANAPC10, ANAPC11, ANAPC13, ANAPC2, ANAPC4, ANAPC5, ANAPC7, ATM, ATR, BUB1, BUB1B, BUB3, CCNA1, CCNA2, CCNB1, CCNB2, CCNB3, CCND1, CCND2, CCND3, CCNE1, CCNE2, CCNH, CDC14A, CDC14B, CDC16,* |

| | |
|---|---|
| | *CDC20, CDC23, CDC25A, CDC25B, CDC25C, CDC26, CDC27, CDC45, CDC6, CDC7, CDK1, CDK2, CDK4, CDK6, CDK7, CDKN1A, CDKN1B, CDKN1C, CDKN2A, CDKN2B, CDKN2C, CDKN2D, CHEK1, CHEK2, CREBBP, CUL1, DBF4, E2F1, E2F2, E2F3, E2F4, E2F5, EP300, ESPL1, FZR1, GADD45A, GADD45B, GADD45G, GSK3B, HDAC1, HDAC2, MAD1L1, MAD2L1, MAD2L2, MCM2, MCM3, MCM4, MCM5, MCM6, MCM7, MDM2, MYC, ORC1, ORC2, ORC3, ORC4, ORC5, ORC6, PCNA, PKMYT1, PLK1, PRKDC, PTTG1, PTTG2, RAD21, RB1, RBL1, RBL2, RBX1, SFN, SKP1, SKP2, SMAD2, SMAD3, SMAD4, SMC1A, SMC1B, SMC3, STAG1, STAG2, TFDP1, TFDP2, TGFB1, TGFB2, TGFB3, TP53, TTK, WEE1, WEE2, YWHAB, YWHAE, YWHAG, YWHAH, YWHAQ, YWHAZ, ZBTB17* |
| p53 | *ATM, CHEK2, ATR, CHEK1, GORAB, CDKN2A, MDM2, MDM4, TP53, CDKN1A, CCND1, CCND2, CCND3, CDK4, CDK6, CCNE1, CCNE2, CDK2, SFN, RPRM, CCNB1, CCNB2, CDK1, GADD45A, GADD45B, GADD45G, GTSE1, FAS, PIDD1, CASP8, BID, BAX, PMAIP1, BBC3, TP53AIP1, SIVA1, BCL2L1, BCL2, TP53I3, EI24, SHISA5, PERP, ZMAT3, SIAH1, CYCS, APAF1, CASP9, CASP3, AIFM2, IGFBP3, IGF1, SERPINE1, ADGRB1, CD82, THBS1, SERPINB5, DDB2, RRM2B, RRM2, SESN1, SESN3, SESN2, PTEN, TSC2, STEAP3, COP1, RCHY1, CCNG1, CCNG2, PPM1D, TP73, TNFRSF10B* |
| Notch | *ADAM17, APH1A, APH1B, ATXN1, ATXN1L, CIR1, CREBBP, CTBP1, CTBP2, DLL1, DLL3, DLL4, DTX1, DTX2, DTX3, DTX3L, DTX4, DVL1, DVL2, DVL3, EP300, HDAC1, HDAC2, HES1, HES5, HEY1, HEY2, HEYL, JAG1, JAG2, KAT2A, KAT2B, LFNG, MAML1, MAML2, MAML3, MFNG, NCOR2, NCSTN, NOTCH1, NOTCH2, NOTCH3, NOTCH4, NUMB, NUMBL, PSEN1, PSEN2, PSENEN, PTCRA, RBPJ, RBPJL, RFNG, SNW1* |
| mTOR | *AKT1, AKT1S1, AKT2, AKT3, ATP6V1A, ATP6V1B1, ATP6V1B2, ATP6V1C1, ATP6V1C2, ATP6V1D, ATP6V1E1, ATP6V1E2, ATP6V1F, ATP6V1G1, ATP6V1G2, ATP6V1G3, ATP6V1H, BRAF, CAB39, CAB39L, CASTOR1, CASTOR2, CHUK, CLIP1, DDIT4, DEPDC5, DEPTOR, DVL1, DVL2, DVL3, EIF4B, EIF4E, EIF4E1B, EIF4E2, EIF4EBP1, FLCN, FNIP1, FNIP2, FZD1, FZD10, FZD2, FZD3, FZD4, FZD5, FZD6, FZD7, FZD8, FZD9, GRB10, GRB2, GSK3B, HRAS, IGF1, IGF1R, IKBKB, INS, INSR, IRS1, KRAS, LAMTOR1, LAMTOR2, LAMTOR3, LAMTOR4, LAMTOR5, LPIN1, LRP5, LRP6, MAP2K1, MAP2K2, MAPK1, MAPK3, MAPKAP1, MIOS, MLST8, MTOR, NPRL2, NPRL3, NRAS, PDPK1, PIK3CA, PIK3CB, PIK3CD, PIK3R1, PIK3R2,* |

| | |
|---|---|
| | *PIK3R3, PRKAA1, PRKAA2, PRKCA, PRKCB, PRKCG, PRR5, PTEN, RAF1, RHEB, RHOA, RICTOR, RNF152, RPS6, RPS6KA1, RPS6KA2, RPS6KA3, RPS6KA6, RPS6KB1, RPS6KB2, RPTOR, RRAGA, RRAGB, RRAGC, RRAGD, SEC13, SEH1L, SESN2, SGK1, SKP2, SLC38A9, SLC3A2, SLC7A5, SOS1, SOS2, STK11, STRADA, STRADB, TBC1D7, TBC1D7-LOC100130357, TELO2, TNF, TNFRSF1A, TSC1, TSC2, TTI1, ULK1, ULK2, WDR24, WDR59, WNT1, WNT10A, WNT10B, WNT11, WNT16, WNT2, WNT2B, WNT3, WNT3A, WNT4, WNT5A, WNT5B, WNT6, WNT7A, WNT7B, WNT8A, WNT8B, WNT9A, WNT9B* |
| Antigen processing and presentation | *IFNG, TNF, PSME1, PSME2, PSME3, HSPA8, HSPA1A, HSPA2, HSPA1L, HSPA1B, HSPA6, HSPA4, HSP90AA1, HSP90AB1, HLA-A, HLA-B, HLA-C, HLA-F, HLA-G, HLA-E, HSPA5, CANX, B2M, PDIA3, CALR, TAPBP, TAP1, TAP2, CD8A, CD8B, CD8B2, KIR3DL2, KIR3DL1, KIR3DL3, KIR2DL2, KIR2DL1, KIR2DL3, KIR2DL4, KIR2DL5A, KLRC1, KLRC2, KLRC3, KLRC4, KLRD1, KIR2DS1, KIR2DS3, KIR2DS4, KIR2DS5, KIR2DS2, IFI30, LGMN, CTSB, HLA-DMA, HLA-DMB, HLA-DOA, HLA-DOB, HLA-DPA1, HLA-DPB1, HLA-DQA1, HLA-DQA2, HLA-DQB1, HLA-DRA, HLA-DRB1, HLA-DRB3, HLA-DRB4, HLA-DRB5, CD74, CTSL, CTSS, CD4, CIITA, RFX5, RFXANK, RFXAP, CREB1, NFYA, NFYB, NFYC* |
| PD-L1 expression and PD-1 checkpoint pathway in cancer | *HIF1A, EGF, EGFR, HRAS, KRAS, NRAS, RAF1, MAP2K1, MAP2K2, MAPK1, MAPK3, FOS, JUN, EML4, ALK, PIK3R1, PIK3R2, PIK3R3, PIK3CA, PIK3CD, PIK3CB, PTEN, AKT1, AKT2, AKT3, MTOR, RPS6KB1, RPS6KB2, CHUK, IKBKB, IKBKG, NFKBIA, NFKBIB, NFKBIE, NFKB1, RELA, IFNG, IFNGR1, IFNGR2, JAK1, JAK2, STAT1, STAT3, TLR2, TLR4, TLR9, TIRAP, MYD88, TRAF6, NFATC1, NFATC2, NFATC3, TICAM1, TICAM2, CD274, PDCD1, PTPN6, PTPN11, BATF3, BATF, BATF2, CSNK2A1, CSNK2A2, CSNK2A3, CSNK2B, CD4, LCK, CD3E, CD3G, CD247, CD3D, ZAP70, MAP3K3, MAP2K3, MAP2K6, MAPK11, MAPK12, MAPK13, MAPK14, LAT, PLCG1, PPP3CA, PPP3CB, PPP3CC, PPP3R1, PPP3R2, RASGRP1, CD28, PRKCQ* |
